# Genomic analysis identifies *Campylobacter concisus* genomospecies 2 as a novel species and proposes the name *Campylobacter oralis*

**DOI:** 10.1101/2025.06.24.661227

**Authors:** Fang Liu, Siying Chen, Joanna M Biazik, Christopher Luk, Anjaneyaswamy Ravipati, Ruiting Lan, Stephen M Riordan, Li Zhang

## Abstract

*Campylobacter concisus* comprises two distinct genomospecies (GS) based on core genome analysis. Here we conducted further analysis and demonstrated that *C. concisus* GS2 strains belong to a novel *Campylobacter* species.

A total of 245 *C. concisus* genomes including 85 GS1 and 160 GS2 strains were analysed. DNA-DNA hybridization (DDH) values between strains of GS1 and GS2 *C. concisus* were from 42.8% to 66%. Average Nucleotide Identity (ANI) between GS1 and GS2 *C. concisus* strains were from 87.8% to 89.7%. The average GC contents of GS1 and GS2 *C. concisus* strains were 37.3% and 39.2% respectively. GS1 and GS2 specific genes have been identified. GS1 and GS2 *C. concisus* strains can also be differentiated based on core genome, 23S rRNA polymorphisms and protein profiles. Their morphology appears different under certain conditions.

In conclusion, ANI and DDH values clearly support that *C. concisus* GS2 as a novel *Campylobacter* species, for which we propose the name *Campylobacter oralis.* The identification of *C. concisus* and *C. oralis* specific genes, along with differences in their protein profiles, morphology, core genome and 23S rRNA phylogeny, further support this classification.

## INTRODUCTION

*Campylobacter concisus* was first isolated by Tanner et al. from human dental specimen in 1981, with strain ATCC 33237 as the type strain [1]. Later studies isolated *C. concisus* from fecal samples and biopsies of patients with gastritis, inflammatory bowel disease (IBD) which includes Crohn’s disease and ulcerative colitis, and other diseases [2–8]. In 2016, Chung et al. conducted a comparative genome analysis of 36 *C. concisus* strains and reported that *C. concisus* contained two GS based on the core genome and identified GS specific genes [9]. Based on the core genome, the *C. concisus* reference strain ATCC33237 is a GS1 strain [9]. Kirk et al. in 2018 examined 113 *C. concisus* genomes and reported that the genomes of GS2 *C. concisus* strains contained higher numbers of genes compared to genomes of GS1 *C. concisus* strains [10]. In 2020, Liu et al. analysed 16 complete sequenced *C. concisus* genomes and reported that the genomes of GS2 *C. concisus* strains had larger sizes and higher GC contents than the genomes of GS1 *C. concisus* strains [11]. In 2021, Cornelius et al. analysed 190 *C. concisus* genomes, also confirmed the existence of GS1 and GS2 *C. concisus* [12].

We hypothesize that GS2 *C. concisus* is a novel species and conducted further analysis of 245 *C. concisus* genomes and also investigated methods that may phenotypically differentiate *C. concisus* GS1 and GS2 strains. Our data support that the GS2 *C. concisus* is a novel *Campylobacter* species, and we propose the name *Campylobacter oralis*. We also examined the presence of *csep1* gene, a molecular marker that is associated with active Crohn’s disease and found that one of these markers, *csep1* gene, was only present in *C. oralis,* not in *C. concisus*.

## MATERIALS AND METHODS

### Analysis of genomes of *C. concisus* GS1 and GS2 strains

The genomes of 245 *C. concisus* strains, including 85 GS1 and 160 GS2 strains, were obtained from National Center for Biotechnology Information (NCBI) genome database for analysis (Supplementary Table 1).

### Phylogenetic analysis based on core genome, 23S and 16S rRNA genes

The core genome of the 245 *C. concisus* strains was generated by using Roary (v3.12.0) without splitting paralogs [13]. The phylogenetic tree of core genome was generated by using FastTree (v2.1.10) based on the approximate-maximum-likelihood method [14]. The sequences of 16S and 23S rRNA gene were used to generate phylogenetic trees based on the maximum-likelihood method (100 bootstrap) by using MEGAcc (v10.1.6), and *Campylobacter jejuni* NCTC113651 was used as an outgroup [15].

### DNA-DNA hybridization (DDH), average nucleotide identity (ANI) and GC content

*In silico* DDH and ANI analyses were performed on all 245 *C. concisus* genomes. DDH values were calculated on Genome-to-Genome Distance Calculator (GGDC 2.1 https://ggdc.dsmz.de). ANI values were generated using fastANI (v1.3) [16, 17]. The GC contents were determined using SeqKit [18].

### Identification of genomospecies-specific genes

The results on the presence and absence of genes among 245 *C. concisus* strains obtained from Roary were further analysed through Scoary (v1.6.16), GS-specific genes were then identified [19]. Redundant genes were excluded until mutually exclusive GS-specific genes were yielded. The presence of each of the candidate GS specific genes in 245 *C. concisus* strains was confirmed by blasting each gene against whole genomes and annotated genes using nucleotide BLAST, and annotated proteins using protein BLAST. In this study, GS specific genes are defined as genes present in GS1 strains but absent in all GS2 strains, *vice versa*.

### Matrix assisted laser desorption ionization-time of flight mass spectrometry (MALDI-TOF MS) analysis

A total of 20 *C. concisus* strains (10 GS1 strains and 10 GS2 strains) were used for this analysis (Supplementary Table 1). The complete genomes of these strains were sequenced previously [11]. These strains were cultured on horse blood agar (HBA) plates under anaerobic conditions supplemented with 5% H_2_ for 48 hours as described previously [20].

Bacterial proteins were prepared using ethanol-formic acid extraction procedure. Briefly, a loopful bacteria of each strain was suspended in 300 µL of distilled water, and 900 µL of ethanol was added. After mixing thoroughly, the bacterial cell suspension was centrifuged at 13,000 × g for 2 mins and the supernatant was discarded. The residual ethanol was removed by repeating centrifugation. The pellet was dried at room temperature for 5 mins and suspended in 5 to 50 µL of formic acid-water (70:30 [vol/vol]), and an equal volume of ethanol was added. Following centrifugation at 13,000 × g for 2 mins, 1 µL of supernatant was spotted on ground steel MALDI target plate and allowed to dry at room temperature prior to overlaying 1 µL of matrix solution.

MALDI-TOF MS analysis was performed using the linear positive ion mode on Ultraflex TOF-TOF mass spectrometer (Bruker Daltonik GmbH, Bremen, Germany). External calibrations were performed using The Bruker Bacterial Test Standard (Bruker Daltanics) according to the manufacturer’s instructions. The raw spectra data were exported to Biotyper Version 4.0 (Build 19) for bacterial identification and FlexAnalysis Version 3.4 for further downstream data analysis.

### Scanning electron microscope (SEM) imaging of GS1 and GS2 *C. concisus* strains

Six *C. concisus* strains were randomly chosen for SEM examination, including three GS1 strains including P26UCO-S2, P10CDO-S2, and P3UCO1; and three GS2 strains including BEO2, P15UCO-S2, and P2CDO4.

*C. concisus* strains were cultured on HBA plates for 24 hours, followed by cultivation in heart infusion broth containing 5% fetal bovine serum for 12 hours. Bacterial cells were collected and fixed overnight at 4 °C in 2.5% glutaraldehyde in 0.2 M sodium phosphate buffer. Fixed samples were dehydrated using a graded series of ethanol washes (30%, 50%, 70%, 80%, 90% and 2 × 100%) for 5 mins in each solution change. Samples were then air-dried using hexamethyldisilazane evaporative dehydration. Samples were then mounted onto SEM stubs, platinum coated and viewed using a FEI Nova NanoSEM 230 (Oregon, USA) operating at 5kV.

### Examination of the presence of *csep1* gene, plasmids pSma1 and pICON in GS1 and GS2 *C. concisus* strains

The *C. concisus csep1* gene, plasmids pSma1 and pICON were previously found to be associated with IBD, including both CD and UC [11, 21]. Their presence in the genomes of 85 GS1 and 160 GS2 strains, as well as in other *Campylobacter* species was examined by NCBI nucleotide BLAST [22].

### Growth characters of GS2 *C. concisus* strains and catalase, oxidase and urease tests

Growth of 10 GS2 *C. concisus* strains with complete sequenced genomes (Supplementary Table 1) under different conditions, including incubation temperature, atmospheric conditions, resistance to bile were examined. Gram staining was performed. Catalase, oxidase and urease tests were also performed.

## RESULTS

### *C. concisus* GS1 and GS2 strains can be separated based on core genome and 23S rRNA gene

The core genome of 245 *C. concisus* strains contained 675 genes, as determined by Roary default setting (the gene is present in more than or equal to 99% of strains). The 85 GS1 *C. concisus* strains and 160 GS2 *C. concisus* strains were clearly separated based on the sequences of core genome (Figure 1). The GS2 *C. concisus* strains contained two clusters, most of the strains belonged to cluster 1. The 23S rRNA gene also consistently separated GS1 and GS2 *C. concisus* strains (Figure 1). However, the sequences of 16S rRNA genes were unable to differentiate GS1 and GS2 *C. concisus* (Supplementary Figure 1).

**Figure 1.**
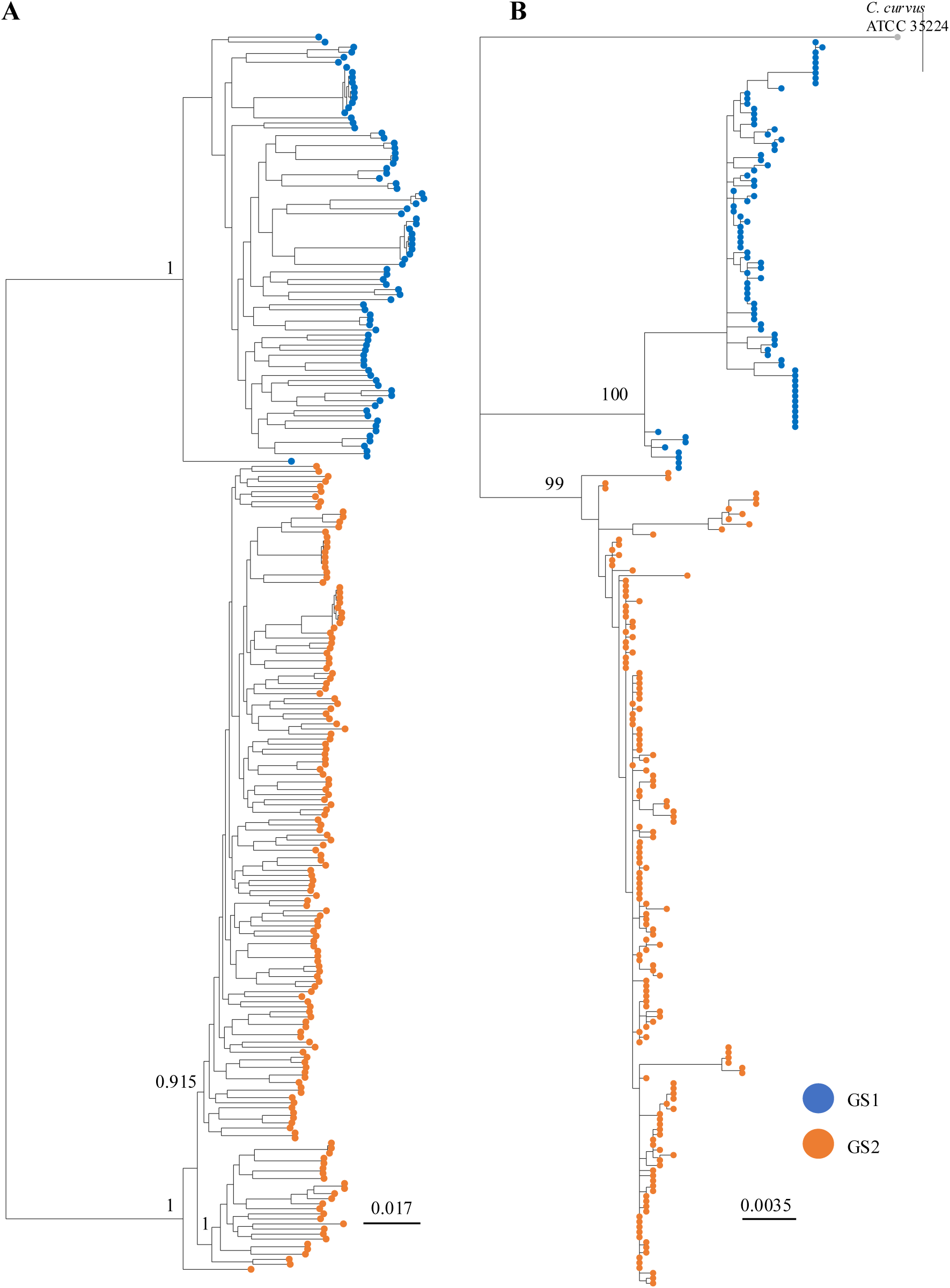
Phylogenetic trees of GS1 and GS2 *C. concisus* based on sequences of *C. concisus* core genome and the 23S rRNA gene. *C. concisus* core genome and 23S rRNA were able to separate GS1 and GS2 strains. **A:** Core genome was generated by Roary (v3.12.0) and the approximate maximum-likelihood tree was generated by FastTree (v2.1.10). **B:** 23S rRNA maximum-likelihood tree (100 bootstrap) was constructed with MEGAXcc. Strains P21CDO-S2 and H25O-S1 were removed for the 23S rRNA phylogenetic tree due to incomplete 23S rRNA sequences.

The average GC content of the 85 GS1 *C. concisus* strains was 37.3%, ranging from 34.7 to 38.2%. The average GC content of the 160 GS2 *C. concisus* strains was 39.2%, ranging from 36.4 to 40.3%.

### The DDH and ANI values between GS1 and GS2 *C. concisus* strains were below the recommended cut off value for defining a bacterial species

The DDH values between each of the 85 GS1 *C. concisus* strains and each of the 160 GS2 *C. concisus* strains were all below 70%. The average DDH value between GS1 and GS2 *C. concisus* strains was 54.9%, ranging from 42.8 to 66% (Table 1).

**Table 1.** DDH and ANI values obtained from analyses of GS1 and GS2 *C. concisus* genomes.

|  |  | **DDH** | **ANI** |
| --- | --- | --- | --- |
| **85 GS1 vs 160 GS2 strains** | Max | 66 | 89.69 |
|  | Min | 42.8 | 87.83 |
|  | Mean | 54.9 | 88.69 |
| **84 GS1 strains versus ATCC 33237** | Max | 99.2 | 99.56 |
|  | Min | 74.5 | 94.01 |
|  | Mean | 82.9 | 94.74 |
| **160 GS2 strains versus ATCC 33237** | Max | 63.7 | 89.15 |
|  | Min | 50.5 | 88.41 |
|  | Mean | 57.4 | 88.73 |

GS: genomospecies. ATCC 33237: *C. concisus* type strain described by Tanner et al [1]

The average ANI value between each of the 85 GS1 *C. concisus* strains and each of the 160 GS2 *C. concisus* strains was 88.7%, ranging from 87.8 to 89.7% (Table 1).

### The DDH values between all GS1 *C. concisus* strains and the *C. concisus* type strain ATCC 33237

The DDH values between the *C. concisus* type strain ATCC 33237 and the 84 remaining GS1 strains were all ≥69% (Table 1). The average DDH values between the 84 GS1 *C. concisus* strains and the *C. concisus* type strain ATCC 33237 was 82.9%, ranging from 74.5 to 99.2 % (Table 1).

The ANI values between the 84 GS1 *C. concisus* strains and the *C. concisus* type strain ATCC 33237 ranged from 93.83 to 99.99% (Table 1).

### The DDH values between all GS2 *C. concisus* strains and the *C. concisus* type strain ATCC 33237 were below 70%

The DDH values between the 160 GS2 strains and the *C. concisus* type strain ATCC 33237 were all below 70%, ranging from 50.5 to 63.7 %. The ANI values between the 160 GS2 *C. concisus* strains and the *C. concisus* type strain ATCC 33237 ranged from 88.41 to 89.15% (Table 1).

### Both GS1 and GS2 *C. concisus* strains carry GS-specific genes

Both GS1 and GS2 *C. concisus* strains contained their respective specific genes. Four GS1 and three GS2 *C. concisus* specific genes were identified (Table 2).

**Table 2.** GS1 and GS2 *C. concisus* specific genes.

| GS | Gene | Annotation | Gene locus | Number of strains present (percentage of strains present) | Protein length (AA) |
| --- | --- | --- | --- | --- | --- |
| **1** | *tehA* | Tellurite resistance protein TehA | CVT06_06385 | 85 (100%) | 319 |
|  | *abgT* | p-aminobenzoyl-glutamate transport protein | CVT06_04695 | 85 (100%) | 512 |
|  | *pstS* | Phosphate-binding protein PstS | CVT06_05910 | 85 (100%) | 290 |
|  |  | Helicase | CVT06_05855 | 81 (95.3%) | 86 |
|  |  | Hypothetical protein | CVT06_06530 | 67 (78.8%) | 57 |
|  |  | Sodium-dependent tyrosine transporter | CVT06_02985 | 65 (76.5%) | 62 |
| **2** | *hcp* | Hydroxylamine reductase | CCC13826_1540 | 160 (100%) | 442 |
|  | *aqpZ* | Aquaporin Z | CCC13826_1636 | 160 (100%) | 236 |
|  |  | Aspartate racemase | CCC13826_1511 | 158 (98.8%) | 230 |
|  | *cas2* | CRISPR-associated endoribonuclease Cas2 | CCC13826_1424 | 121 (75.6%) | 92 |
|  | *hgdC* | Activator of 2-hydroxyglutaryl-CoA dehydratase | CCC13826_1133 | 116 (72%) | 259 |
|  | *dnaX* | DNA polymerase III, gamma and tau subunits | CCC13826_1448 | 115 (71.9%) | 554 |
|  | *cas8* | CRISPR/Cas system-associated protein Cas8, type I-B/HMARI | CCC13826_1419 | 98 (61.3%) | 588 |

GS: genomospecies. AA: amino acid. GS specific genes: present in strains from one GS and absent in all the strains from the other GS.

### MALDI-TOF MS profiles were different between GS1 and GS2 *C. concisus* strains

On MALDI-TOF MS analysis, protein peaks specific to GS1 or GS2 *C. concisus* strains were identified. Two protein peaks, at m/z 7401 and 7557, were found to be present only in GS1 *C. concisus* strains, with the prevalence being 80 and 60% respectively (Table 3). Four protein peaks with m/z 6924, 7424, 8045 and 10359, were present only in GS2 *C. concisus* strains, with the prevalence being 90%, 90%, 50% and 50% respectively (Table 3). Combination of m/z 6924 with any of the other three protein peaks can 100% differentiate GS2 from GS1 strains.

**Table 3.** MALDI-TOF MS markers differentiating GS1 and GS2 *C. concisus*.

| Strain ID | GS | Average m/z |  |  |  |  |  |
| --- | --- | --- | --- | --- | --- | --- | --- |
|  |  | 6924.389 | 7401.524 | 7424.504 | 7557.485 | 8045.274 | 10359.59 |
| P10CDO-S2 | 1 |  | 7392.891 |  |  |  |  |
| P26UCO-S2 | 1 |  | 7409.682 |  | 7558.543 |  |  |
| H25O-S1 | 1 |  |  |  |  |  |  |
| P3UCO1 | 1 |  | 7411.752 |  | 7556.284 |  |  |
| P19CDO-S1 | 1 |  |  |  | 7558.897 |  |  |
| H17O-S1 | 1 |  | 7373.999 |  |  |  |  |
| H28O-S1 | 1 |  | 7406.009 |  |  |  |  |
| H1O1 | 1 |  | 7404.886 |  | 7555.046 |  |  |
| P27CDO-S2 | 1 |  | 7403.849 |  | 7557.604 |  |  |
| P28CDO-S1 | 1 |  | 7409.12 |  | 7558.533 |  |  |
| P1CDO2 | 2 | 6924.038 |  | 7426.252 |  | 8044.699 |  |
| P1CDO3 | 2 | 6926.038 |  |  |  |  | 10359.56 |
| P15UCO-S2 | 2 | 6924.789 |  | 7425.441 |  |  |  |
| P2CDO4 | 2 | 6923.513 |  | 7424.174 |  | 8044.692 | 10359.5 |
| H14O-S1 | 2 |  |  | 7424.693 |  | 8045.149 | 10358.19 |
| P16UCO-S2 | 2 | 6923.745 |  | 7426.193 |  | 8045.902 | 10360.75 |
| H19O-S1 | 2 | 6924.265 |  | 7422.043 |  |  | 10359.98 |
| P20CDO-S2 | 2 | 6921.875 |  | 7424.294 |  |  |  |
| 13826 | 2 | 6926.661 |  | 7423.399 |  | 8045.93 |  |
| P13UCO-S1 | 2 | 6924.579 |  | 7424.045 |  |  |  |

GS: genomospecies

### GS1 and GS2 *C. concisus* strains showed difference in morphology

Electron microscopy examination showed that strains of GS1 and GS2 *C. concisus* differ in morphology. GS1 *C. concisus* strains were straight to curved rods and the shapes of GS2 *C. concisus* strains were spiral to curved (Figure 2).

**Figure 2.**
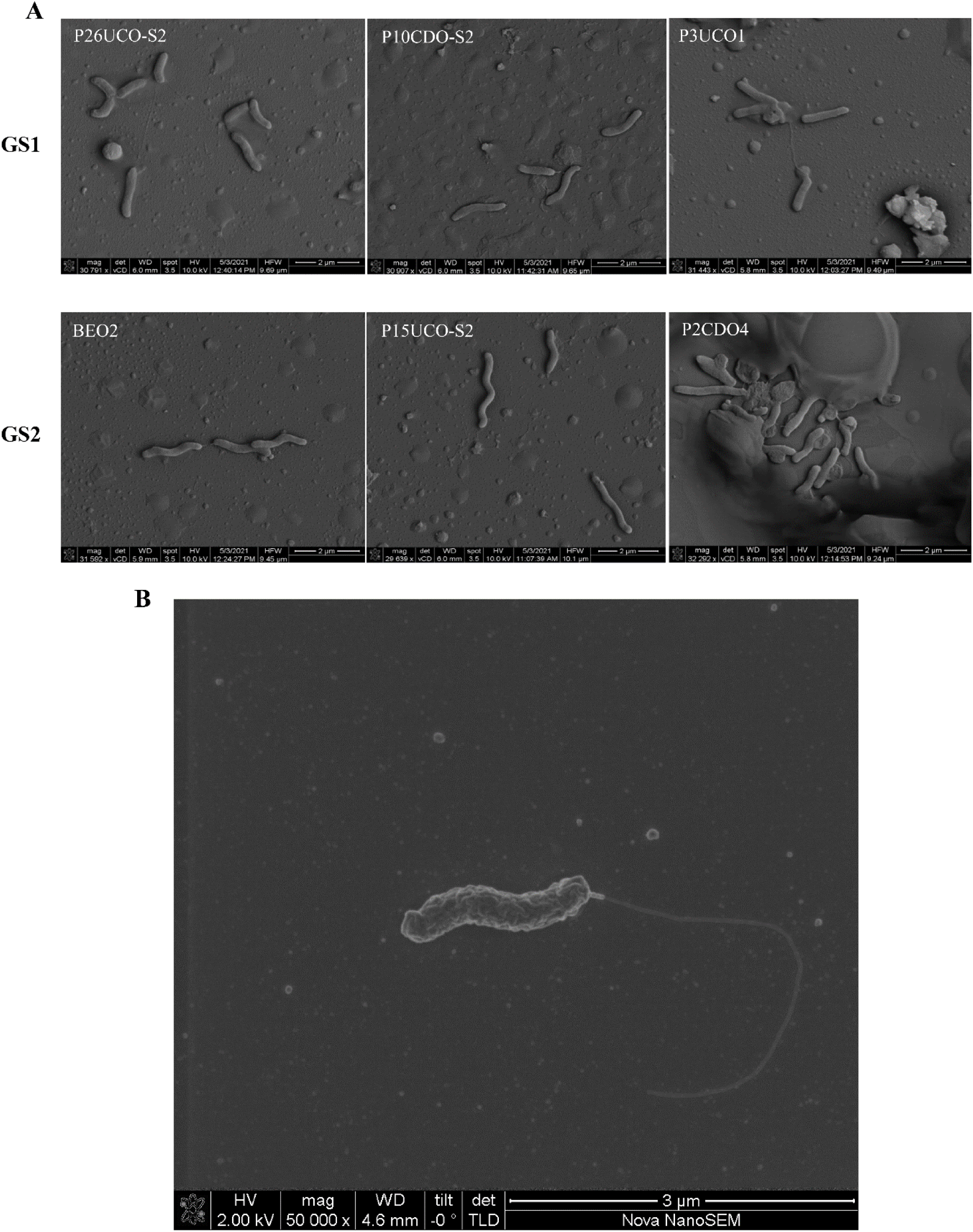
Scanning electron microscope images showing the different morphologies of GS1 and GS2 *C. concisus.* A: *C. concisus* strains were cultured on HBA plates for 24 hours followed by cultivation in heart infusion broth with 5% fetal bovine serum for 12 hours. Bacterial cells were then subjected to scanning electron microscopic examination. Top row: GS1 strains including P26UCO-S2, P10CDO-S2 and P3UCO1. Bottom row: GS2 strains including BEO2, P15UCO-S2 and P2CDO4. Magnification: 30000×, scale bars represent 2 µm. **B:** Showing the single polar flagellum of *C. oralis*. Magnification:500000×. GS: genomospecies.

### The Crohn’s disease associated *csep1* gene and plasmids were predominately present in GS2 *C. concisus* strains

The Crohn’s disease associated *csep1* gene was found in the chromosome of 38.75% (62/160) GS2 *C. concisus* strains and none of the GS1 strains. The Crohn’s disease associated pICON plasmid was found in two GS1 strains (2/85, 2.4%), four GS2 strains (4/160, 2.5%) and one strain of *Campylobacter showae* (B32_SW) [23]. The severe ulcerative colitis associated pSma1 plasmid was found in three GS1 strains (3/85, 3.5%) and 13 GS2 strains (13/160, 8.1%) (Supplementary Table 1).

## DISCUSSION

We previously reported that *C. concisus* consisted of GS1 and GS2 strains based on the difference in the core genome, which was confirmed by other researchers [9–12]. Our findings in this study show that GS2 *C. concisus* is a novel *Campylobacter* species, we propose naming it *C. oralis,* whereas GS1 *C. concisus* solely remains as *C. concisus*.

The recommended cut off DDH and ANI values for defining bacterial species are 70% DDH and 95% ANI [17, 24–26]. Both the DDH and ANI values between *C. oralis* and *C. concisus* strains were clearly below these criteria (Table 1). Phenotypically, *C. oralis* has a spiral to curved rod shape and *C. concisus* has a straight or curved rod shape under the culture conditions that we have used (Figure 2). MALDI-TOF MS analysis identified *C. oralis* and *C. concisus* specific protein peaks. Our study and the study by On et al. consistently identified four *C. oralis* protein peaks with the m/z 6924, 7424, 8045 and 10359 (Table 3) [27]. Combination of these protein peaks can 100% differentiate *C. oralis* from *C. concisus*. Based on our genome analysis, *C. oralis* and *C. concisus* specific genes were not related to common metabolic pathways, making biochemical differentiation difficult. PCR targeting 23S rRNA gene can also be used as a quick method for differentiating *C. oralis* from *C. concisus* and the sequences of PCR primers were reported in previous studies [28, 29]. The 16S rRNA gene cannot differentiate *C. oralis* from *C. concisus* (Supplementary Figure 1).

We and other researchers previously reported the association between *C. concisus* and IBD [2–5]. In these earlier studies, the identification of *C. concisus* was based on 16S rRNA gene, thus both *C. oralis* and *C. concisus* were detected. Recently, we also reported the associations between *csep1* gene, pICON and pSma1 plasmids with IBD [11, 21]. In this study, we found that Crohn’s disease associated *csep1* gene was present in 38.75% of *C. oralis* strains and none of the *C. concisus* strains. The plasmid pICON is rare, we found it in only seven strains of three *Campylobacter* species (four *C. oralis* strains, two *C. concisus* strains and one *C. showae* strain), interestingly all these strains were isolated from patients with Crohn’s disease [10, 21, 23]. The severe ulcerative colitis associated pSma1 plasmid was found in less than 5% of *C. concisus* strains and 10% of *C. oralis* strains. These suggest that *C. oralis* may play a more important role in the development of Crohn’s disease than *C. concisus*.

### Description of *C. oralis*

*C. oralis* bacterial cells are Gram negative, with a curved to spiral shape, 0.5 – 0.8 × 4 – 8 µm, having rounded ends, motile with darting motility by means of a single polar flagellum (Figure 2). Colonies are circular, with convex elevation, translucent, smooth margins, no pigmentation, and no observable haemolysis on HBA plates. The growth and biochemical characteristics of *C. oralis* are listed in Table 4.

**Table 4.**
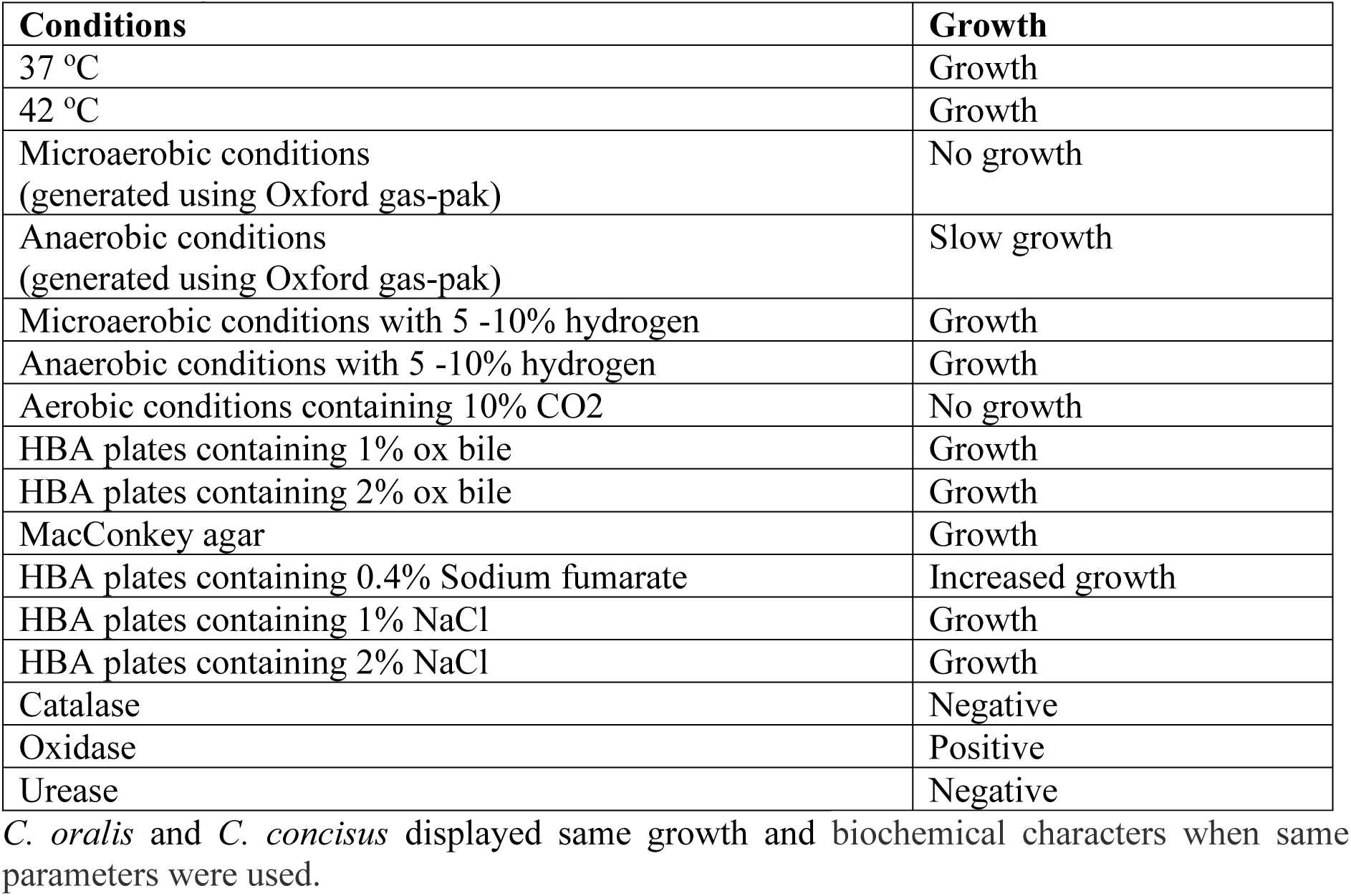
The growth and biochemical characters of *C. oralis*.

| Conditions | Growth |
| --- | --- |
| 37 °C | Growth |
| 42 °C | Growth |
| Microaerobic conditions<br>(generated using Oxford gas-pak) | No growth |
| Anaerobic conditions<br>(generated using Oxford gas-pak) | Slow growth |
| Microaerobic conditions with 5 -10% hydrogen | Growth |
| Anaerobic conditions with 5 -10% hydrogen | Growth |
| Aerobic conditions containing 10% CO<sub>2</sub> | No growth |
| HBA plates containing 1% ox bile | Growth |
| HBA plates containing 2% ox bile | Growth |
| MacConkey agar | Growth |
| HBA plates containing 0.4% Sodium fumarate | Increased growth |
| HBA plates containing 1% NaCl | Growth |
| HBA plates containing 2% NaCl | Growth |
| Catalase | Negative |
| Oxidase | Positive |
| Urease | Negative |

*C. oralis* and *C. concisus* displayed same growth and biochemical characters when same parameters were used.

P15UCO-S2 strain is the type strain of *C. oralis*. This strain was isolated from saliva of a patient with ulcerative colitis and its complete genome has been sequenced [11]. The DDH values between the 159 *C. oralis* strains and the *C. oralis* type strain P15UCO-S2 ranged from 66.2 to 98.7%. The ANI values between the 159 *C. oralis* strains and the type strain P15UCO-S2 ranged from 93.6 to 98.7%. Based on these analyses, we recommend 66.2% DDH and 93.6% ANI with the type strain P15UCO-S2 as the cut off values for identifying *C. oralis*. These values are able to exclude *C. concisus* strains, the DDH and ANI values between the 85 *C. concisus* strains and the *C. oralis* type strain P15UCO-S2 were 53.4 – 64.9% and 88.2 – 89.3% respectively. The *C. oralis* type strain has been deposited in the DSMZ under the number DSM 115209.

*C. oralis* was most closely related to *C. concisus* (Supplementary Table 2). *C. oralis* and *C. concisus* can be differentiated based on the core genome sequence, 23S rRNA gene, DDH value, ANI value and species-specific genes. *C. oralis* and *C. concisus* show different morphology under electron microscope under certain culture conditions. PCR targeting 23S rRNA gene can be used to differentiate *C. oralis* and *C. concisus* in clinical laboratories. MALDI-TOF MS is a method that can be potentially used to differentiate *C. oralis* and *C. concisus*.

## AUTHOR CONTRIBUTION

Fang Liu: Conducted analysis of *csep1* gene and plasmids. Performed confirmation analysis on GS specific genes.

Siying Chen: Performed genome analysis, most of the experiments and prepared the type strain for submission to ATCC.

Joanna M Biazik: Generated SEM pictures for Figure 2A using bacterial strains cultured by Siying Chen.

Christopher Luk: Conducted catalase, oxidase and urease tests.

Anjaneyaswamy Ravipati: Generated data for Table 3 using bacterial strains cultured by Siying Chen.

Ruiting Lan: Provided critical discussion during this project.

Li Zhang and Stephen M Riordan: Conceived the project and Li Zhang supervised the project.

Li Zhang, Fang Liu and Siying Chen played a major role in writing the manuscript. All authors have read the manuscript and provided feedback.

## ACKNOWLEDGEMENT

Picture in Figure 2B was generated by Vikneswari Mahendran during her study for a Masters by Research degree in Associate Professor Li Zhang group at the University of New South Wales in 2011.

