## Supplementary Material for "Genomic analysis identifies *Campylobacter concisus* genomospecies 2 as a novel species and proposes the name *Campylobacter oralis*"

**Supplementary Table 1. GS1 and GS2 *C. concisus* strains used in this study.**

| Strain ID | GS | NCBI accession | Isolation source | Isolation country | Isolation year | Number of contigs | N50 | Ref |
|---|---|---|---|---|---|---|---|---|
| 2009-158448 | 1 | GCA_902460545.1 | Faeces | Denmark | Unknown | 127 | 26966 | [1] |
| 2009-173039 | 1 | GCA_902460475.1 | Faeces | Denmark | Unknown | 53 | 59456 | [1] |
| 2009-42653 | 1 | GCA_902460445.1 | Faeces | Denmark | Unknown | 44 | 65473 | [1] |
| 2010-112100-F | 1 | GCA_902460415.1 | Faeces | Denmark | Unknown | 49 | 67761 | [1] |
| 2010-113332-F | 1 | GCA_902460785.1 | Faeces | Denmark | Unknown | 101 | 39586 | [1] |
| 2010-164712 | 1 | GCA_902460565.1 | Faeces | Denmark | Unknown | 91 | 39715 | [1] |
| 2010-1718 | 1 | GCA_902460625.1 | Faeces | Denmark | Unknown | 107 | 31608 | [1] |
| 2010-25654-F | 1 | GCA_902460665.1 | Faeces | Denmark | Unknown | 39 | 81434 | [1] |
| 2010-25654-O | 1 | GCA_902460735.1 | Saliva | Denmark | Unknown | 79 | 37302 | [1] |
| 2010-347972 | 1 | GCA_902460585.1 | Faeces | Denmark | Unknown | 175 | 29064 | [1] |
| 2010-378007-F | 1 | GCA_902460685.1 | Faeces | Denmark | Unknown | 71 | 47212 | [1] |
| 2010-378007-O | 1 | GCA_902460705.1 | Saliva | Denmark | Unknown | 29 | 112662 | [1] |
| 2010-6073 | 1 | GCA_902460885.1 | Faeces | Denmark | Unknown | 95 | 33564 | [1] |
| 2010-8194 | 1 | GCA_902460805.1 | Faeces | Denmark | Unknown | 100 | 30171 | [1] |
| 2012-164712 | 1 | GCA_902460845.1 | Faeces | Denmark | Unknown | 37 | 174192 | [1] |
| 2012-37302 | 1 | GCA_902460855.1 | Faeces | Denmark | Unknown | 125 | 25704 | [1] |
| 2013-87946 | 1 | GCA_902460875.1 | Faeces | Denmark | Unknown | 112 | 27923 | [1] |
| AAUH-16UCf2 | 1 | GCA_002914365.1 | Faeces | Denmark | 2016 | 138 | 21709 | [2] |
| AAUH-12CDti2-a | 1 | GCA_002913715.1 | Intestinal biopsy | Denmark | 2016 | 11 | 315326 | [2] |
| AAUH-12CDdes3 | 1 | GCA_002914355.1 | Intestinal biopsy | Denmark | 2016 | 73 | 49375 | [2] |
| AAUH-5CDo | 1 | GCA_002913685.1 | Saliva | Denmark | 2016 | 260 | 89584 | [2] |
| AAUH-11UCdes-a*^ | 1 | GCA_002913155.1 | Intestinal biopsy | Denmark | 2016 | 62 | 58169 | [2] |
| AAUH-11UCo^ | 1 | GCA_002914195.1 | Saliva | Denmark | 2016 | 149 | 161715 | [2] |
| AAUH-37UCo-a | 1 | GCA_002913185.1 | Saliva | Denmark | 2016 | 54 | 89957 | [2] |

| | | | | | | | | |
|---|---|---|---|---|---|---|---|---|
| AAUH-16UCf[#] | 1 | GCA_002914165.1 | Faeces | Denmark | 2016 | 78 | 392237 | [2] |
| AAUH-12CDsig | 1 | GCA_002913145.1 | Intestinal biopsy | Denmark | 2016 | 42 | 257808 | [2] |
| AAUH-16UCo-a[#] | 1 | GCA_002913095.1 | Saliva | Denmark | 2016 | 60 | 151248 | [2] |
| AAUH-16UCf3 | 1 | GCA_002914145.1 | Faeces | Denmark | 2016 | 66 | 64374 | [2] |
| AAUH-12CDce | 1 | GCA_002914115.1 | Intestinal biopsy | Denmark | 2016 | 44 | 483873 | [2] |
| AAUH-12CDdes4 | 1 | GCA_002914105.1 | Intestinal biopsy | Denmark | 2016 | 38 | 158715 | [2] |
| AAUH-12CDdes2 | 1 | GCA_002913085.1 | Intestinal biopsy | Denmark | 2016 | 55 | 158794 | [2] |
| AAUH-12CDtra2-a | 1 | GCA_002914065.1 | Intestinal biopsy | Denmark | 2016 | 41 | 266692 | [2] |
| AAUH-12CDti5-a | 1 | GCA_002913065.1 | Intestinal biopsy | Denmark | 2016 | 37 | 187882 | [2] |
| AAUH-20UCo[#] | 1 | GCA_002913045.1 | Saliva | Denmark | 2016 | 47 | 235464 | [2] |
| AAUH-12CDti4-a | 1 | GCA_002912995.1 | Intestinal biopsy | Denmark | 2016 | 54 | 83693 | [2] |
| AAUH-12CDo | 1 | GCA_002913015.1 | Saliva | Denmark | 2016 | 43 | 90391 | [2] |
| AAUH-6HCo-a | 1 | GCA_002912985.1 | Saliva | Denmark | 2016 | 91 | 34235 | [2] |
| AAUH-8HCo | 1 | GCA_002912965.1 | Saliva | Denmark | 2016 | 148 | 23759 | [2] |
| AAUH-9HCasc | 1 | GCA_002914025.1 | Intestinal biopsy | Denmark | 2016 | 89 | 41150 | [2] |
| AAUH-8HCo-a | 1 | GCA_002912935.1 | Saliva | Denmark | 2016 | 137 | 25859 | [2] |
| AAUH-14HCce | 1 | GCA_002912925.1 | Intestinal biopsy | Denmark | 2016 | 161 | 22276 | [2] |
| AAUH-2010376221 | 1 | GCA_002912895.1 | Faeces | Denmark | 2016 | 84 | 48012 | [2] |
| AAUH-55UCtra-a | 1 | GCA_002912885.1 | Intestinal biopsy | Denmark | 2016 | 91 | 37875 | [2] |
| AAUH-48UCo-a | 1 | GCA_002914005.1 | Saliva | Denmark | 2016 | 87 | 45066 | [2] |
| AAUH-51UCf | 1 | GCA_002912785.1 | Faeces | Denmark | 2016 | 66 | 100202 | [2] |
| AAUH-11HCf | 1 | GCA_002913965.1 | Faeces | Denmark | 2016 | 235 | 14154 | [2] |
| AAUH-2012179281 | 1 | GCA_002913945.1 | Faeces | Denmark | 2016 | 39 | 138913 | [2] |
| AAUH-11HCo-a | 1 | GCA_002912715.1 | Saliva | Denmark | 2016 | 237 | 14621 | [2] |
| AAUH-3HCo | 1 | GCA_002912395.1 | Saliva | Denmark | 2016 | 572 | 26143 | [2] |
| P10CDO-S1 | 1 | GCA_003049705.1 | Saliva | Australia | Unknown | 14 | 275885 | [3] |
| P10CDO-S2[&] | 1 | GCA_003048695.2 | Saliva | Australia | Unknown | 1 | 1932636 | [3] |

| | | | | | | | | |
|---|---|---|---|---|---|---|---|---|
| P28CDO-S1& | 1 | GCA_003048405.1 | Saliva | Australia | Unknown | 10 | 1034549 | [3] |
| ATCC 33237 | 1 | GCA_001298465.1 | Oral cavity | USA | 1974 | 1 | 1840041 | NA |
| ATCC 51562 | 1 | GCA_000466745.1 | Faeces | UK | Unknown | 21 | 361423 | NA |
| AUS22-Bd2 | 1 | GCA_002092845.1 | Intestinal biopsy | Australia | 2013 | 42 | 125724 | [4] |
| H12O-S1 | 1 | GCA_003048755.1 | Saliva | Australia | Unknown | 11 | 687684 | [3] |
| H27O-S1 | 1 | GCA_003048905.1 | Saliva | Australia | Unknown | 10 | 1209207 | [3] |
| H34O-S1 | 1 | GCA_015229985.1 | Saliva | Australia | 2019 | 14 | 296998 | [3] |
| H10O-S1 | 1 | GCA_003048575.1 | Saliva | Australia | Unknown | 13 | 267337 | [3] |
| H21O-S3 | 1 | SRX2253589 | Saliva | Australia | Unknown | 25 | 369626 | [3] |
| H17O-S1& | 1 | SRX2253584 | Saliva | Australia | Unknown | 15 | 986534 | [3] |
| H30O-S1 | 1 | GCA_003048925.1 | Saliva | Australia | Unknown | 31 | 98274 | [3] |
| H35O-S1 | 1 | GCA_015229955.1 | Saliva | Australia | 2019 | 18 | 256719 | [3] |
| P19CDO-S1& | 1 | GCA_003049105.1 | Saliva | Australia | Unknown | 9 | 1174594 | [3] |
| H15O-S1 | 1 | GCA_003048995.1 | Saliva | Australia | Unknown | 14 | 194730 | [3] |
| P26UCO-S2& | 1 | GCA_003048595.2 | Saliva | Australia | Unknown | 2 | 1894099 | [3] |
| P27CDO-S2& | 1 | GCA_003048775.2 | Saliva | Australia | Unknown | 2 | 1831320 | [3] |
| H28O-S1& | 1 | GCA_003048495.1 | Saliva | Australia | Unknown | 33 | 91334 | [3] |
| H28O-S2 | 1 | GCA_003049735.1 | Saliva | Australia | Unknown | 14 | 360820 | [3] |
| Lasto205.94 | 1 | GCA_002165895.1 | Faeces | South Africa | 1994 | 39 | 179898 | [5] |
| Lasto220.96 | 1 | GCA_002165875.1 | Faeces | South Africa | 1996 | 13 | 349534 | [5] |
| Lasto28.99 | 1 | GCA_002165785.1 | Faeces | South Africa | 1999 | 30 | 150974 | [5] |
| Lasto393.96 | 1 | GCA_002165855.1 | Faeces | South Africa | 1996 | 15 | 256243 | [5] |
| Lasto61.99 | 1 | GCA_002114545.1 | Faeces | South Africa | 1999 | 18 | 260338 | [5] |
| Lasto64.99 | 1 | GCA_002165775.1 | Faeces | South Africa | 1999 | 20 | 190490 | [5] |
| H1O1& | 1 | GCA_015679965.1 | Saliva | Australia | 2009 | 1 | 1861371 | [3] |
| RCH 26 | 1 | GCA_002092835.1 | Faeces | Australia | 2008 | 23 | 300266 | [4] |

| | | | | | | | | |
|---|---|---|---|---|---|---|---|---|
| RMIT-JF1 | 1 | GCA_001891085.1 | Oral cavity | Australia | 2013 | 25 | 338852 | [4] |
| P3UCB1 | 1 | GCA_015680005.1 | Intestinal biopsy | Australia | 2009 | 1 | 1831655 | [6] |
| P3UCO1[&] | 1 | GCA_015679985.1 | Saliva | Australia | 2009 | 1 | 1800519 | [3] |
| H26O-S1 | 1 | GCA_003048965.1 | Saliva | Australia | Unknown | 6 | 1025414 | [3] |
| H24O-S1 | 1 | GCA_003048555.1 | Saliva | Australia | Unknown | 88 | 39412 | [3] |
| H25O-S1[&] | 1 | GCA_003048535.1 | Saliva | Australia | Unknown | 22 | 180040 | [3] |
| P20CDO-S4 | 1 | SRX2253579 | Saliva | Australia | Unknown | 10 | 1298069 | [3] |
| P25CDO-S3 | 1 | GCA_003049085.1 | Saliva | Australia | Unknown | 18 | 273413 | [3] |
| 13826[&] | 2 | GCA_000017725.2 | Faeces | Denmark | Unknown | 3 | 2052007 | [1] |
| 2009-118452 | 2 | GCA_902460495.1 | Faeces | Denmark | Unknown | 146 | 26714 | [1] |
| 2009-119100* | 2 | GCA_902460395.1 | Faeces | Denmark | Unknown | 131 | 27698 | [1] |
| 2009-129008* | 2 | GCA_902460485.1 | Faeces | Denmark | Unknown | 199 | 19347 | [1] |
| 2009-130586* | 2 | GCA_902460425.1 | Faeces | Denmark | Unknown | 150 | 23125 | [1] |
| 2009-75710 | 2 | GCA_902460405.1 | Faeces | Denmark | Unknown | 99 | 34189 | [1] |
| 2009-75775* | 2 | GCA_902460465.1 | Faeces | Denmark | Unknown | 108 | 72398 | [1] |
| 2009-86120 | 2 | GCA_902460525.1 | Faeces | Denmark | Unknown | 120 | 32258 | [1] |
| 2009-91522 | 2 | GCA_902460515.1 | Faeces | Denmark | Unknown | 75 | 58036 | [1] |
| 2010-112100-O | 2 | GCA_902460505.1 | Saliva | Denmark | Unknown | 33 | 183901 | [1] |
| 2010-112708 | 2 | GCA_902460455.1 | Faeces | Denmark | Unknown | 185 | 21026 | [1] |
| 2010-112758 | 2 | GCA_902460435.1 | Faeces | Denmark | Unknown | 77 | 104654 | [1] |
| 2010-112825* | 2 | GCA_902460535.1 | Faeces | Denmark | Unknown | 51 | 73883 | [1] |
| 2010-113332-O* | 2 | GCA_902460715.1 | Saliva | Denmark | Unknown | 39 | 115430 | [1] |
| 2010-113862 | 2 | GCA_902460605.1 | Faeces | Denmark | Unknown | 140 | 30832 | [1] |
| 2010-113862-O | 2 | GCA_902460725.1 | Saliva | Denmark | Unknown | 58 | 73494 | [1] |
| 2010-115605-F* | 2 | GCA_902460595.1 | Faeces | Denmark | Unknown | 147 | 30412 | [1] |
| 2010-131105* | 2 | GCA_902460675.1 | Faeces | Denmark | Unknown | 34 | 154245 | [1] |
| 2010-16206* | 2 | GCA_902460635.1 | Faeces | Denmark | Unknown | 54 | 101107 | [1] |

| | | | | | | | | |
|---|---|---|---|---|---|---|---|---|
| 2010-30795 | 2 | GCA_902460555.1 | Faeces | Denmark | Unknown | 146 | 36383 | [1] |
| 2010-30800 | 2 | GCA_902460645.1 | Faeces | Denmark | Unknown | 143 | 27804 | [1] |
| 2010-31374 | 2 | GCA_902460615.1 | Faeces | Denmark | Unknown | 69 | 59551 | [1] |
| 2010-33561*^ | 2 | GCA_902460695.1 | Faeces | Denmark | Unknown | 53 | 101406 | [1] |
| 2010-34330* | 2 | GCA_902460575.1 | Faeces | Denmark | Unknown | 62 | 87626 | [1] |
| 2010-36743* | 2 | GCA_902460745.1 | Faeces | Denmark | Unknown | 54 | 86581 | [1] |
| 2010-43100* | 2 | GCA_902460895.1 | Faeces | Denmark | Unknown | 113 | 81275 | [1] |
| 2010-88823 | 2 | GCA_902460775.1 | Faeces | Denmark | Unknown | 139 | 28977 | [1] |
| 2012-191940 | 2 | GCA_902460835.1 | Faeces | Denmark | Unknown | 171 | 20870 | [1] |
| 2013-101463 | 2 | GCA_902460865.1 | Faeces | Denmark | Unknown | 81 | 56878 | [1] |
| 2013-39845* | 2 | GCA_902460905.1 | Faeces | Denmark | Unknown | 154 | 29360 | [1] |
| 2013-42088 | 2 | GCA_902460755.1 | Faeces | Denmark | Unknown | 55 | 76427 | [1] |
| AAUH-22UCpp-a* | 2 | GCA_002913785.1 | Intestinal biopsy | Denmark | 2016 | 51 | 108686 | [2] |
| AAUH-10UCil-a* | 2 | GCA_002913765.1 | Intestinal biopsy | Denmark | 2016 | 29 | 197033 | [2] |
| AAUH-20UCf# | 2 | GCA_002913745.1 | Faeces | Denmark | 2016 | 81 | 45189 | [2] |
| AAUH-40UCf | 2 | GCA_002913705.1 | Faeces | Denmark | 2016 | 182 | 20799 | [2] |
| AAUH-15UCdp-a# | 2 | GCA_002913665.1 | Intestinal biopsy | Denmark | 2016 | 203 | 52907 | [2] |
| AAUH-10UCf2* | 2 | GCA_002914345.1 | Faeces | Denmark | 2016 | 55 | 180736 | [2] |
| AAUH-43UCce-a* | 2 | GCA_002913635.1 | Intestinal biopsy | Denmark | 2016 | 206 | 73289 | [2] |
| AAUH-16UCdp5# | 2 | GCA_002914325.1 | Intestinal biopsy | Denmark | 2016 | 58 | 142542 | [2] |
| AAUH-15UCdp# | 2 | GCA_002913605.1 | Intestinal biopsy | Denmark | 2016 | 130 | 35022 | [2] |
| AAUH-37UCf | 2 | GCA_002914295.1 | Faeces | Denmark | 2016 | 97 | 87543 | [2] |
| AAUH-8UCpp# | 2 | GCA_002913575.1 | Intestinal biopsy | Denmark | 2016 | 101 | 39071 | [2] |
| AAUH-8UCpp-a# | 2 | GCA_002913075.1 | Intestinal biopsy | Denmark | 2016 | 101 | 39071 | [2] |
| AAUH-39CDrec-a | 2 | GCA_002913565.1 | Intestinal biopsy | Denmark | 2016 | 103 | 40955 | [2] |
| AAUH-11UCsig-a | 2 | GCA_002913525.1 | Intestinal biopsy | Denmark | 2016 | 46 | 159029 | [2] |

| | | | | | | | | |
|---|---|---|---|---|---|---|---|---|
| AAUH-16UCdp3 | 2 | GCA_002913505.1 | Intestinal biopsy | Denmark | 2016 | 78 | 64884 | [2] |
| AAUH-8UCo*# | 2 | GCA_002913465.1 | Saliva | Denmark | 2016 | 52 | 124439 | [2] |
| AAUH-4UCti-a | 2 | GCA_002914285.1 | Intestinal biopsy | Denmark | 2016 | 77 | 97971 | [2] |
| AAUH-4UCti | 2 | GCA_002913485.1 | Intestinal biopsy | Denmark | 2016 | 76 | 92998 | [2] |
| AAUH-7UCil | 2 | GCA_002914265.1 | Intestinal biopsy | Denmark | 2016 | 80 | 108467 | [2] |
| AAUH-35UCdp | 2 | GCA_002913445.1 | Intestinal biopsy | Denmark | 2016 | 139 | 127580 | [2] |
| AAUH-12CDtra-a | 2 | GCA_002913425.1 | Intestinal biopsy | Denmark | 2016 | 73 | 63299 | [2] |
| AAUH-35UCf | 2 | GCA_002913395.1 | Faeces | Denmark | 2016 | 95 | 87016 | [2] |
| AAUH-43UCf* | 2 | GCA_002913385.1 | Faeces | Denmark | 2016 | 69 | 62023 | [2] |
| AAUH-39CDf | 2 | GCA_002914245.1 | Faeces | Denmark | 2016 | 74 | 136855 | [2] |
| AAUH-3UCce | 2 | GCA_002913365.1 | Intestinal biopsy | Denmark | 2016 | 96 | 103045 | [2] |
| AAUH-35UCil4-a | 2 | GCA_002914225.1 | Intestinal biopsy | Denmark | 2016 | 83 | 80285 | [2] |
| AAUH-35UCil-a | 2 | GCA_002913345.1 | Intestinal biopsy | Denmark | 2016 | 75 | 97085 | [2] |
| AAUH-39CDti-a | 2 | GCA_002914185.1 | Intestinal biopsy | Denmark | 2016 | 76 | 120353 | [2] |
| AAUH-35UCpp | 2 | GCA_002913325.1 | Intestinal biopsy | Denmark | 2016 | 47 | 134251 | [2] |
| AAUH-3UCce2 | 2 | GCA_002913305.1 | Intestinal biopsy | Denmark | 2016 | 57 | 101107 | [2] |
| AAUH-16UCdp# | 2 | GCA_002913275.1 | Intestinal biopsy | Denmark | 2016 | 84 | 162832 | [2] |
| AAUH-35UCil2-a | 2 | GCA_002913245.1 | Intestinal biopsy | Denmark | 2016 | 53 | 138370 | [2] |
| AAUH-35UCil3-a | 2 | GCA_002913265.1 | Intestinal biopsy | Denmark | 2016 | 45 | 97130 | [2] |
| AAUH-12CDrec-a* | 2 | GCA_002913225.1 | Intestinal biopsy | Denmark | 2016 | 74 | 102065 | [2] |
| AAUH-15UCpp# | 2 | GCA_002913195.1 | Intestinal biopsy | Denmark | 2016 | 56 | 162832 | [2] |
| AAUH-9UCdp | 2 | GCA_002914085.1 | Intestinal biopsy | Denmark | 2016 | 54 | 81490 | [2] |
| AAUH-25Df* | 2 | GCA_002914045.1 | Faeces | Denmark | 2016 | 57 | 87543 | [2] |
| AAUH-1Dasc | 2 | GCA_002912865.1 | Intestinal biopsy | Denmark | 2016 | 241 | 13132 | [2] |
| AAUH-10HCdes4 | 2 | GCA_002912805.1 | Intestinal biopsy | Denmark | 2016 | 98 | 45360 | [2] |
| AAUH-10HCdes2 | 2 | GCA_002912825.1 | Intestinal biopsy | Denmark | 2016 | 84 | 59463 | [2] |
| AAUH-10HCdes7 | 2 | GCA_002912815.1 | Intestinal biopsy | Denmark | 2016 | 77 | 51438 | [2] |

| | | | | | | | | |
|---|---|---|---|---|---|---|---|---|
| AAUH-1Dtra | 2 | GCA_002913985.1 | Intestinal biopsy | Denmark | 2016 | 93 | 39458 | [2] |
| AAUH-10HCdes5 | 2 | GCA_002912745.1 | Intestinal biopsy | Denmark | 2016 | 85 | 39333 | [2] |
| AAUH-10HCdes3 | 2 | GCA_002912735.1 | Intestinal biopsy | Denmark | 2016 | 66 | 74769 | [2] |
| AAUH-10HCdes | 2 | GCA_002912705.1 | Intestinal biopsy | Denmark | 2016 | 56 | 65110 | [2] |
| AAUH-47UCil-a* | 2 | GCA_002912675.1 | Intestinal biopsy | Denmark | 2016 | 93 | 42174 | [2] |
| AAUH-59UCpp-a* | 2 | GCA_002912665.1 | Intestinal biopsy | Denmark | 2016 | 78 | 102305 | [2] |
| AAUH-10HCce | 2 | GCA_002912615.1 | Intestinal biopsy | Denmark | 2016 | 90 | 43541 | [2] |
| AAUH-15HCti* | 2 | GCA_002913905.1 | Intestinal biopsy | Denmark | 2016 | 129 | 24816 | [2] |
| AAUH-48UCdp-a | 2 | GCA_002913895.1 | Intestinal biopsy | Denmark | 2016 | 212 | 19218 | [2] |
| AAUH-10HCtra | 2 | GCA_002912585.1 | Intestinal biopsy | Denmark | 2016 | 75 | 60348 | [2] |
| AAUH-47UCil* | 2 | GCA_002912595.1 | Intestinal biopsy | Denmark | 2016 | 73 | 70435 | [2] |
| AAUH-12HCf# | 2 | GCA_002913885.1 | Faeces | Denmark | 2016 | 167 | 22427 | [2] |
| AAUH-48UCil-a | 2 | GCA_002912605.1 | Intestinal biopsy | Denmark | 2016 | 61 | 65623 | [2] |
| AAUH-10HCdes6 | 2 | GCA_002912565.1 | Intestinal biopsy | Denmark | 2016 | 110 | 36601 | [2] |
| AAUH-9HCce* | 2 | GCA_002912525.1 | Intestinal biopsy | Denmark | 2016 | 192 | 21463 | [2] |
| AAUH-19HCf2* | 2 | GCA_002913865.1 | Faeces | Denmark | 2016 | 157 | 23859 | [2] |
| AAUH-2HCtra* | 2 | GCA_002912505.1 | Intestinal biopsy | Denmark | 2016 | 97 | 52482 | [2] |
| AAUH-19HCf* | 2 | GCA_002913835.1 | Faeces | Denmark | 2016 | 85 | 48338 | [2] |
| AAUH-58UCo | 2 | GCA_002912515.1 | Saliva | Denmark | 2016 | 108 | 34563 | [2] |
| AAUH-44UCsig6* | 2 | GCA_002912485.1 | Intestinal biopsy | Denmark | 2016 | 255 | 16657 | [2] |
| AAUH-20HCrec-a | 2 | GCA_002912465.1 | Intestinal biopsy | Denmark | 2016 | 129 | 84768 | [2] |
| AAUH-20HCsig-a | 2 | GCA_002912425.1 | Intestinal biopsy | Denmark | 2016 | 197 | 33006 | [2] |
| AAUH-20HCasc | 2 | GCA_002913825.1 | Intestinal biopsy | Denmark | 2016 | 443 | 19626 | [2] |
| AAUH-49UCpp-a* | 2 | GCA_002913805.1 | Intestinal biopsy | Denmark | 2016 | 304 | 52807 | [2] |
| AAUH-49UCil-a* | 2 | GCA_002912365.1 | Intestinal biopsy | Denmark | 2016 | 255 | 41082 | [2] |
| AAUH-3HCce2* | 2 | GCA_002912385.1 | Intestinal biopsy | Denmark | 2016 | 553 | 31996 | [2] |
| AAUH-49UCf* | 2 | GCA_002912335.1 | Faeces | Denmark | 2016 | 161 | 97148 | [2] |

| | | | | | | | | |
|---|---|---|---|---|---|---|---|---|
| AAUH-1Dce-a | 2 | GCA_002912325.1 | Intestinal biopsy | Denmark | 2016 | 584 | 45493 | [2] |
| P21CDO-S1 | 2 | SRX2253580 | Saliva | Australia | Unknown | 32 | 215002 | [3] |
| P21CDO-S2*# | 2 | SRX2253577 | Saliva | Australia | Unknown | 68 | 332372 | [3] |
| P21CDO-S4 | 2 | SRX2253578 | Saliva | Australia | Unknown | 38 | 129907 | [3] |
| P1CDO2& | 2 | GCA_003048675.2 | Saliva | Australia | Unknown | 1 | 2031332 | [3] |
| P1CDO3*& | 2 | GCA_003048685.2 | Saliva | Australia | Unknown | 1 | 2051359 | [3] |
| P2CDO3* | 2 | SRX2253601 | Saliva | Australia | Unknown | 72 | 277481 | [3] |
| P2CDO4*^& | 2 | GCA_003048375.1 | Saliva | Australia | 2009 | 2 | 1975443 | [3] |
| P2CDO-S6* | 2 | SRX2253599 | Saliva | Australia | Unknown | 53 | 278451 | [3] |
| P6CDO1 | 2 | GCA_003048505.1 | Saliva | Australia | Unknown | 21 | 348922 | [3] |
| P11CDO-S1* | 2 | GCA_003048875.2 | Saliva | Australia | Unknown | 2 | 2025227 | [3] |
| H32O-S1 | 2 | GCA_015230035.1 | Saliva | Australia | 2019 | 29 | 151581 | [3] |
| H32O-S2 | 2 | GCA_015230015.1 | Saliva | Australia | 2019 | 39 | 108327 | [3] |
| ATCC 51561 | 2 | GCA_000466705.1 | Faeces | Sweden | Unknown | 69 | 111029 | NA |
| H33O-S1 | 2 | GCA_015229965.1 | Saliva | Australia | 2019 | 54 | 75419 | [3] |
| B124_Slimy-large | 2 | GCA_902460825.1 | Intestinal biopsy | UK | Unknown | 285 | 17284 | [1] |
| B124_Slimy-small | 2 | GCA_902460815.1 | Intestinal biopsy | UK | Unknown | 39 | 220370 | [1] |
| B124_Small-clear | 2 | GCA_902460795.1 | Intestinal biopsy | UK | Unknown | 125 | 38449 | [1] |
| B124_Small-grey | 2 | GCA_902460765.1 | Intestinal biopsy | UK | Unknown | 58 | 120359 | [1] |
| B38_Tiny-mucoid | 2 | GCA_902460915.1 | Intestinal biopsy | UK | Unknown | 46 | 75981 | [1] |
| H29O-S1* | 2 | GCA_003048705.1 | Saliva | Australia | Unknown | 47 | 76358 | [3] |
| CCUG 19995* | 2 | GCA_002165815.1 | Faeces | Sweden | 1987 | 44 | 215794 | [5] |
| H21O-S1* | 2 | SRX2253591 | Saliva | Australia | Unknown | 38 | 112460 | [3] |
| H21O-S2 | 2 | SRX2253590 | Saliva | Australia | Unknown | 40 | 138316 | [3] |
| H21O-S5 | 2 | SRX2253588 | Saliva | Australia | Unknown | 73 | 75277 | [3] |
| H22O-S1 | 2 | SRX2253593 | Saliva | Australia | Unknown | 76 | 162003 | [3] |

| | | | | | | | | |
|---|---|---|---|---|---|---|---|---|
| H14O-S1[&] | 2 | SRX2253585 | Saliva | Australia | Unknown | 33 | 322175 | [3] |
| P7UCO-S2* | 2 | GCA_003048895.1 | Saliva | Australia | Unknown | 26 | 226478 | [3] |
| P26UCO-S1* | 2 | GCA_003048445.1 | Saliva | Australia | Unknown | 41 | 122849 | [3] |
| P16UCO-S2*[&] | 2 | GCA_003048645.1 | Saliva | Australia | Unknown | 11 | 341483 | [3] |
| P27CDO-S1* | 2 | GCA_003048765.1 | Saliva | Australia | Unknown | 30 | 205442 | [3] |
| H16O-S1 | 2 | GCA_003048615.2 | Saliva | Australia | Unknown | 1 | 1987364 | [3] |
| H9O-S1*[#] | 2 | GCA_003048815.1 | Saliva | Australia | 2015 | 19 | 203101 | [3] |
| H9O-S2* | 2 | GCA_015680025.1 | Saliva | Australia | 2015 | 2 | 2025058 | [3] |
| P24CDO-S2 | 2 | SRX2253582 | Saliva | Australia | Unknown | 26 | 184230 | [3] |
| P24CDO-S3 | 2 | SRX2253583 | Saliva | Australia | Unknown | 57 | 85614 | [3] |
| P24CDO-S4 | 2 | SRX2253581 | Saliva | Australia | Unknown | 40 | 136251 | [3] |
| Lasto127.99 | 2 | GCA_002165825.1 | Faeces | South Africa | 1999 | 44 | 134605 | [5] |
| P15UCO-S2[#&] | 2 | GCA_003048845.1 | Saliva | Australia | 2015 | 24 | 1943962 | [3] |
| H19O-S1[&] | 2 | GCA_003116505.1 | Saliva | Australia | Unknown | 32 | 197598 | [3] |
| H11O-S1 | 2 | GCA_003049025.1 | Saliva | Australia | Unknown | 27 | 128646 | [3] |
| H11O-S2* | 2 | GCA_003048605.1 | Saliva | Australia | Unknown | 18 | 200644 | [3] |
| H31O-S1 | 2 | GCA_015230075.1 | Saliva | Australia | 2019 | 27 | 165853 | [3] |
| H7O-S1 | 2 | GCA_003049045.1 | Saliva | Australia | Unknown | 17 | 346163 | [3] |
| H36O-S1 | 2 | GCA_015229935.1 | Saliva | Australia | 2019 | 35 | 219194 | [3] |
| H23O-S1 | 2 | SRX2253592 | Saliva | Australia | Unknown | 33 | 298329 | [3] |
| P13UCO-S1*[&] | 2 | GCA_003048835.2 | Saliva | Australia | Unknown | 2 | 1998513 | [3] |
| P13UCO-S3* | 2 | GCA_015679945.1 | Saliva | Australia | Unknown | 3 | 214312 | [3] |
| P18CDO-S1* | 2 | GCA_003048475.1 | Saliva | Australia | Unknown | 94 | 32966 | [3] |
| UNSW1 | 2 | GCA_000466665.1 | Intestinal biopsy | Australia | Unknown | 72 | 117975 | [7] |
| UNSW2 | 2 | GCA_000466725.1 | Intestinal biopsy | Australia | Unknown | 98 | 89312 | [7] |
| UNSW3 | 2 | GCA_000466645.1 | Intestinal biopsy | Australia | Unknown | 61 | 92608 | [7] |

| | | | | | | | | |
|---|---|---|---|---|---|---|---|---|
| UNSWCD | 2 | GCA_000259315.1 | Intestinal biopsy | Australia | 2008 | 86 | 64047 | [7] |
| UNSWCS* | 2 | GCA_000466685.1 | Faeces | Australia | Unknown | 177 | 68143 | [7] |
| H20O-S1* | 2 | GCA_003048985.1 | Saliva | Australia | Unknown | 26 | 131516 | [3] |
| H3O1* | 2 | GCA_003049065.1 | Saliva | Australia | Unknown | 22 | 211732 | [3] |
| P20CDO-S1* | 2 | SRX2253594 | Saliva | Australia | Unknown | 75 | 227524 | [3] |
| P20CDO-S2*^& | 2 | SRX2253603 | Saliva | Australia | Unknown | 55 | 266446 | [3] |
| P20CDO-S3*^ | 2 | SRX2253602 | Saliva | Australia | Unknown | 71 | 197586 | [3] |
| BEO1* | 2 | GCA_020351935.1 | Saliva | Australia | 2020 | 93 | 37628 | [8] |
| BEO2* | 2 | GCA_020351945.1 | Saliva | Australia | 2020 | 95 | 29126 | [8] |

*Positive for *csep1* gene. ^Positive for pICON plasmid. #Positive for pSma1 plasmid. &Strains used for MALDI-TOF MS. NCBI accession number indicates genome assembly accession number (with prefix GCA) or SRA accession number (with prefix SRX). NA: not available.

**Supplementary Table 2 dDDH and ANI values between *C. oralis* type strain P15UCO-S2 and type strain of other *Campylobacter* species**

| **Strain** | **dDDH (%)** | **ANI (%)** |
|---|---|---|
| *Campylobacter concisus* ATCC 33237 | 63.7 | 88.84 |
| *Campylobacter curvus* ATCC 35224 | 18.2 | 74.23 |
| *Campylobacter anatolicus* faydin-G140 | 14.9 | 72.62 |
| *Campylobacter massiliensis* Marseille-Q3452 | 14.8 | 72.22 |
| *Campylobacter mucosalis* ATCC 43264 | 14.8 | 71.84 |
| *Campylobacter showae* ATCC 51146 | 14.7 | 71.81 |
| *Campylobacter rectus* ATCC 33238 | 14.0 | 70.69 |
| *Campylobacter pinnipediorum* subsp. *pinnipediorum* RM17260 | 13.7 | 69.88 |
| *Campylobacter geochelonis* DSM 102159 | 13.5 | 68.90 |
| *Campylobacter iguaniorum* 1485E | 13.5 | 69.04 |
| *Campylobacter lanienae* NCTC 13004 | 13.5 | 68.06 |
| *Campylobacter armoricus* CCUG 73571 | 13.4 | 66.22 |
| *Campylobacter fetus* subsp. *fetus* NCTC10842 | 13.4 | 67.72 |
| *Campylobacter gracilis* ATCC 33236 | 13.4 | 68.33 |
| *Campylobacter hyointestinalis* subsp. *hyointestinalis* CCUG 14169 | 13.4 | 68.27 |
| *Campylobacter peloridis* LMG 23910 | 13.4 | 66.46 |
| *Campylobacter volucris* LMG 24380 | 13.4 | 66.17 |
| *Campylobacter bilis* VicNov18 | 13.3 | 65.63 |
| *Campylobacter blaseri* LMG 30333 | 13.3 | 67.34 |
| *Campylobacter coli* ATCC 33559 | 13.3 | 66.44 |
| *Campylobacter corcagiensis* LMG 27932 | 13.3 | 67.22 |
| *Campylobacter insulaenigrae* NCTC 12927 | 13.3 | 65.76 |
| *Campylobacter jejuni* subsp. *jejuni* ATCC 33560 | 13.3 | 66.11 |
| *Campylobacter lari* subsp. *lari* ATCC 35221 | 13.3 | 66.36 |
| *Campylobacter novaezeelandiae* B423b | 13.3 | 65.61 |
| *Campylobacter ornithocola* LMG 29815 | 13.3 | 66.28 |
| *Campylobacter portucalensis* FMV-PI01 | 13.3 | 67.29 |

| | | |
|---|---|---|
| *Campylobacter sputorum bv. sputorum* CCUG 9728 | 13.3 | 67.84 |
| *Campylobacter subantarcticus* LMG 24377 | 13.3 | 66.07 |
| *Campylobacter taeniopygiae* MIT10-5678 | 13.3 | 66.09 |
| *Campylobacter ureolyticus* DSM 20703 | 13.3 | 67.16 |
| *Campylobacter aviculae* MIT17-670 | 13.2 | 66.10 |
| *Campylobacter avium* LMG 24591 | 13.2 | 66.22 |
| *Campylobacter cuniculorum* LMG 24588 | 13.2 | 65.67 |
| *Campylobacter estrildidarum* MIT17-664 | 13.2 | 66.05 |
| *Campylobacter hepaticus* HV10 | 13.2 | 65.45 |
| *Campylobacter hominis* NCTC 13146 | 13.2 | 66.94 |
| *Campylobacter upsaliensis* NCTC 11541 | 13.2 | 66.12 |
| *Campylobacter vulpis* 251/13 | 13.2 | 66.19 |
| *Campylobacter canadensis* LMG 24001 | 13.1 | 65.52 |
| *Campylobacter helveticus* ATCC 51209 | 13.1 | 66.58 |
| *Candidatus Campylobacter infans* 19S00001 | 13.1 | 66.83 |

**Supplementary Figure 1**

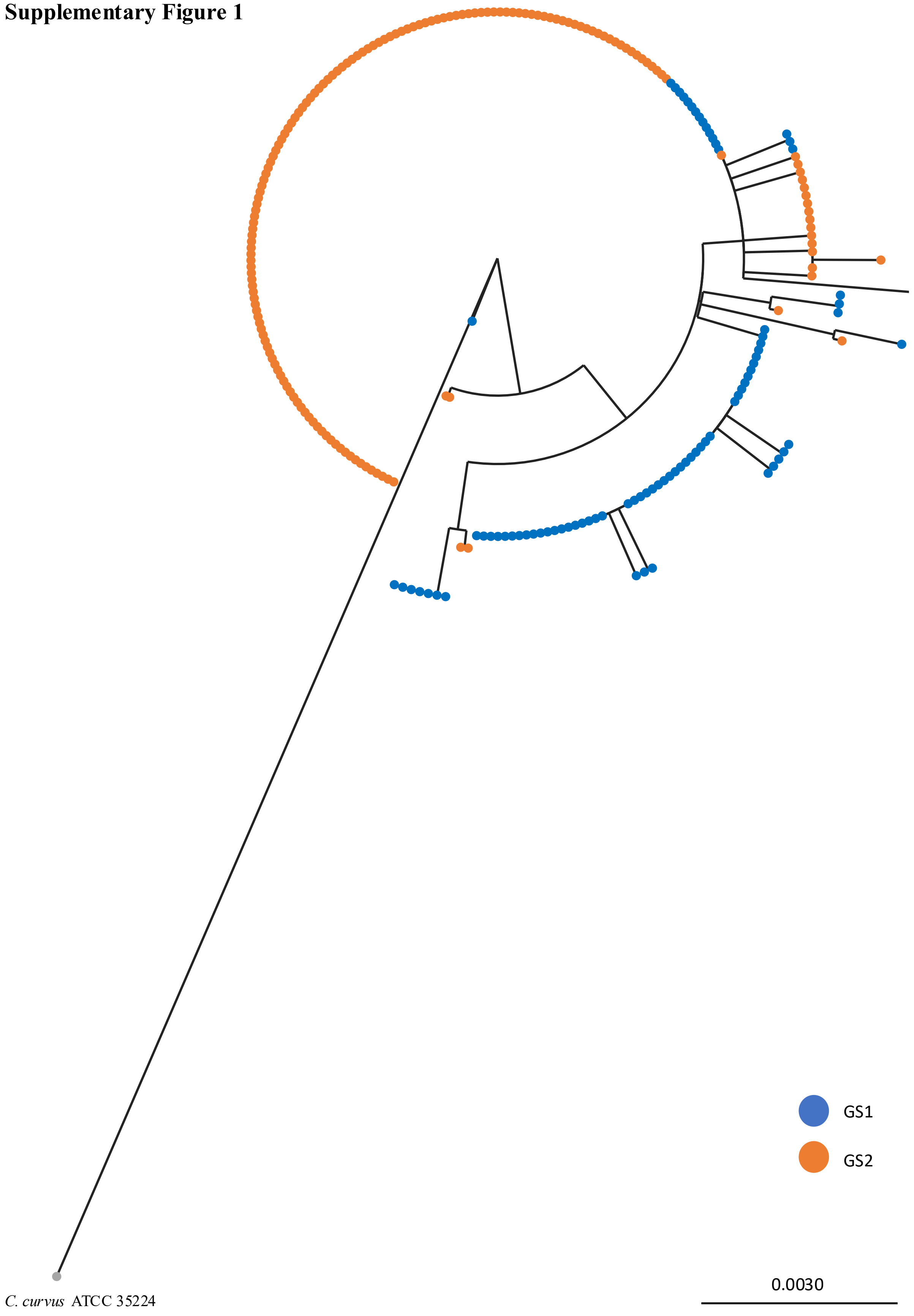


**Supplementary Figure 1 Phylogenetic tree of GS1 and GS2 *C. concisus* based on sequences of 16S rRNA gene.** *C. concisus* 16S rRNA gene was unable to separate GS1 and

GS2 strains. 16S rRNA maximum-likelihood tree (100 bootstrap) was constructed with MEGAXcc.
